# Evolutionary divergence of novel open reading frames in cichlids speciation

**DOI:** 10.1101/2020.03.13.991182

**Authors:** Shraddha Puntambekar, Rachel Newhouse, Jaime San Miguel Navas, Ruchi Chauhan, Grégoire Vernaz, Thomas Willis, Matthew T. Wayland, Yagnesh Urmania, Eric A. Miska, Sudhakaran Prabakaran

**Affiliations:** Department of Biology, Indian Institute of Science Education and Research, Pune, Maharashtra, 411008, India; Department of Genetics, University of Cambridge, Downing Site, CB2 3EH, UK; The Wellcome Trust/CRUK Gurdon Institute, University of Cambridge, Cambridge, CB2 1QN, UK; Wellcome Sanger Institute, Wellcome Genome Campus, Cambridge CB10 1SA, UK; Department of Zoology, University of Cambridge, Downing Site, CB2 3EH, UK; Cambridge Centre for Proteomics, Department of Biochemistry, University of Cambridge, Tennis Court Road, Cambridge, CB2 1QR, United Kingdom; St Edmund’s College, University of Cambridge, CB3 0BN, UK

## Abstract

Novel open reading frames (nORFs) with coding potential may arise from noncoding DNA. Not much is known about their emergence, functional role, fixation in a population or contribution to adaptive radiation. Cichlids fishes exhibit extensive phenotypic diversification and speciation. Encounters with new environments alone are not sufficient to explain this striking diversity of cichlid radiation because other taxa coexistent with the Cichlidae demonstrate lower species richness. Wagner et al analyzed cichlid diversification in 46 African lakes and reported that both extrinsic environmental factors and intrinsic lineage-specific traits related to sexual selection have strongly influenced the cichlid radiation ^1^, which indicates the existence of unknown molecular mechanisms responsible for rapid phenotypic diversification, such as emergence of novel open reading frames (nORFs). In this study, we integrated transcriptomic and proteomic signatures from two tissues of two cichlids species, identified nORFs and performed evolutionary analysis on these nORF regions. Our results suggest that the time scale of speciation of the two species and evolutionary divergence of these nORF genomic regions are similar and indicate a potential role for these nORFs in speciation of the cichlid fishes.

## Introduction

Rapid evolution of the genome, especially the intergenic regions, have been postulated to contribute to genetic diversity ^2 3 4 5 6^. In Drosophila, yeasts, stickleback fishes, and even in humans some intergenic regions have been shown to evolve into ‘de novo’ or ‘orphan’ or ‘proto’ genes, and they were more often shown to be specifically expressed as transcripts in tissues associated with male reproduction ^5,7,8^, suggesting a sexual or gamete selection. Subsequently, de novo genes were identified in many other organisms ^9, 10^. In this work, we address these de novo genes, and other as yet uncharacterized open reading frames, such as alternate open reading frames, short open reading frames, stop codon read throughs, intron insertions and so on, as described by Prabakaran et al, 2014 ^11^, as novel open reading frames (nORFs) because the definition of de novo gene is stringent in that it must have a monophyletic distribution in one focal clade while being absent from organisms outside this clade.

Not much is known about how the transition from intergenic regions to nORFs to expressed nORFs to protein coding nORFs occurs ^12^. nORFs can emerge from pre-existing genes, for example, through gene duplications that can have an adaptive benefit or other mechanisms such as gene fusion or fission, horizontal gene transfer, exon shuffling and retroposition ^13 10^. nORFs can also emerge ‘de novo’ as has been shown to have emerged from sequences such as long noncoding RNAs ^14^ or through ‘mixed origin mechanisms’ ^15^, overprinting ^16^-alternative open reading frame transcription ^10 15 16^, intron insertion (exonization) or extension of reading frames ^17 18 19^. Although this mechanism was discounted and dismissed ^20 21^, it is gaining credence as work from our own lab and from that of others show that de novo gene emergence, expression and translation is more pervasive than previously known ^11,22^.

With the advent of ultra-deep sequencing technologies, we are beginning to observe the expression of large intergenic regions ^23^. Results from our own work (manuscript in review) and from others have indicated that nORF transcripts are indeed expressed at a lower abundance compared to already fixed known protein coding transcripts ^3,4^ and their expression is not ‘noisy’ to be dismissed as inconsequential for biological functions. nORF transcripts are shown to become fixed ^24,25^ and we have shown that some nORFs can express noncanonical proteins that can be biologically regulated with potential functions ^11 26^ thus indicating that there might be a selection process. However, some studies, including ours, have demonstrated that nORF encoded proteins have propensity for increased disorder ^27^ than known canonical proteins, hence, it is not clear whether the noncanonical proteins contribute to the fitness of the organisms, or whether their biological activities are effectively neutral without any deleterious consequences. One study has demonstrated that some noncanonical proteins evolve neutrally, and that can at some point they acquire new functions ^28^. Another study has demonstrated that nORFs pervasively emerge from noncoding regions but are rapidly lost again, while only a relatively few are retained much longer ^29^.

The central question as to whether transcription emerged first or whether nORFs emerged first from intergenic regions has not been determined yet. Most studies that have attempted to investigate the emergence of nORFs have used comparative rather than population genetic approaches using just the transcriptomic and genomic data with limited phylogenetic analysis and more importantly have constrained the analyses to genes that have remained fixed over a long time scale. Another limitation of these studies is that they have been conducted mostly using model organisms such as drosophila ^2^ and yeasts ^30,31^ that have not been under natural selection for many generations. For these reasons, we have attempted to investigate the emergence of nORFs from intergenic regions in cichlid fishes - a natural model system to investigate adaptive radiation ^32,33^, using the proteogenomic approach that we developed ^11^ - to identify nORFs, base-wise conservation-acceleration (CONACC) ^34^ analysis - to estimate nonneutral substitution rates of these nORFs, and Bayesian Evolutionary Analysis Sampling Trees model (BEAST) analysis ^35^ - to identify the time-scale divergence of these nORFs. The central questions that we attempted to answer are whether nORFs can emerge from noncoding regions, and if they do, whether their emergence can indicate or provide answers to explain the rapid adaptive radiation of the cichlid fishes. The reason why we chose this model system is explained in depth below.

The family Cichlidae is one of the most species-rich families in vertebrates that has fascinated biologists since Darwin. The primary hotspots of biodiversity for this family are in the East African Great Lakes, namely, Lakes Tanganyika, Malawi and Victoria; together they harbor more than two thousand cichlid species. In each of these lakes, cichlids have evolved independently and vary remarkably in behavior, ecology and morphology. The largest radiations, which in Lakes Victoria, Malawi and Tanganyika, have generated between 250 (Tanganyika) and 500 (Malawi and Victoria) species per lake, took no more than 15,000 to 100,000 years for Victoria and less than 5 million years for Malawi, but 10–12 million years for Lake Tanganyika ^33^.

A genome-wide study of 73 Malawi cichlid species reported a low (0.1-0.25%) average sequence divergence between species pairs, indicating highly similar genomes ^36^. Further, a comparative genomic analysis of three morphologically and ecologically distinct cichlid species from Lake Victoria found highly similar degrees of genetic distance and polymorphism consistent with conservation of protein-coding regions ^37^. A study between two East African cichlid species, *Astatotilapia burtoni* and *Ophthalmotilapia ventralis,* reported a genetic distance of 1.75% when annotated and unannotated transcripts were considered, but only 0.95% when including protein-coding sequences ^32^. The genetic differences responsible for some specific cichlid traits are known; for example, *bmp4*’s influence on jaw morphology ^38,39^, the expression patterns of egg-spots and blotches on fin tissue ^40^, role of gonadotropin-releasing hormone (GnRH) ^41^ and multiple steroid receptors (estrogen, androgen and corticosteroid receptors) ^42^; in chemosensory and auditory plasticity respectively and the divergence in visual pigmentation ‘opsin’ genes affecting mate selection ^43 44^. Sequencing of the genomes of five cichlid species revealed accelerated protein-coding sequence evolution, divergence of regulatory elements, regulation by novel micro-RNAs, and divergence in gene expression associated with transposable element insertions ^33^. Organisms that undergo extensive speciation with diverse phenotypic variation, such as the cichlids, therefore must have highly ‘evolvable’ genomes - genome that is more ‘evolvable’ in the noncoding regions than in the coding regions because genetic, transcriptomic and proteomic diversity among the cichlids appears too low to account for the striking phenotypic diversity of the taxon.

Cichlids and Stickleback fishes are therefore fantastic model systems to investigate the emergence of nORFs in the noncoding regions. Previous work from our lab has revealed functionally active regions in the noncoding genome of many species that have not yet been classified as genes ^11^. We show that these noncoding regions, which include - intron insertions, stop-codon read throughs, upstream insertions, antisense translation, alternate open reading frame translation, intergenic region translation are not only translated but they can form structures and are biologically regulated indicating potential functions. Therefore we embarked on a proteogenomic analysis to identify the emergence of nORFs in two tissues of two cichlid species *Oreochromis niloticus* (Nile tilapia, ON) and *Pundamilia nyererei* (Makobe Island, PN) (**Fig. 1a**) and to investigate whether these nORFs can help explain the speciation of these two species in a short geological time scale. The entire workflow is illustrated in **Fig. 1b**. These species are genetically similar but phenotypically divergent. PN is a rock-dwelling lacustrine fish ^45^, whilst ON dwells in rivers ^46^. They differ also in diet, ON is an omnivore with a primarily plant-based diet and PN is a carnivore ^47, 48^. ON has a more plain colouration whereas PN males have yellow flanks and red dorsal regions, a trait that is subject to sexual selection ^45,49^. We compared the expression of transcripts between the species in two metabolically-active tissues, the testes and liver. We chose to study testes because it is known that the highest number and highest expression levels of de novo genes has been observed in testes of drosophila and humans, which indicates that they may contribute to unique traits ^7,8^. We chose liver under the rationale that the distinct diets of the species might be accompanied by divergent liver transcriptomes. Analysis of transcript expression in the testes allowed comparison of extent of divergence in sex and non-sex traits.

**Figure 1:**
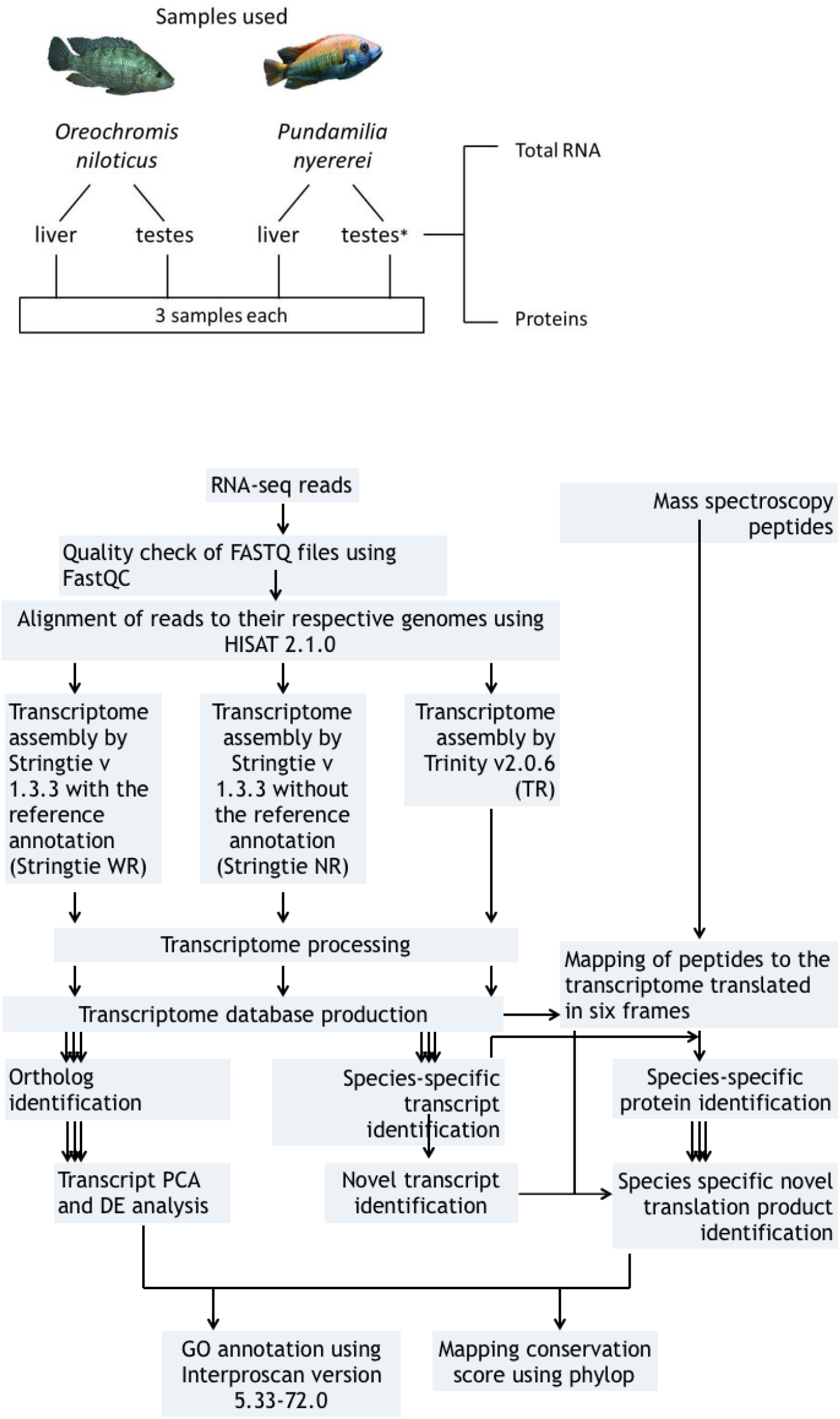
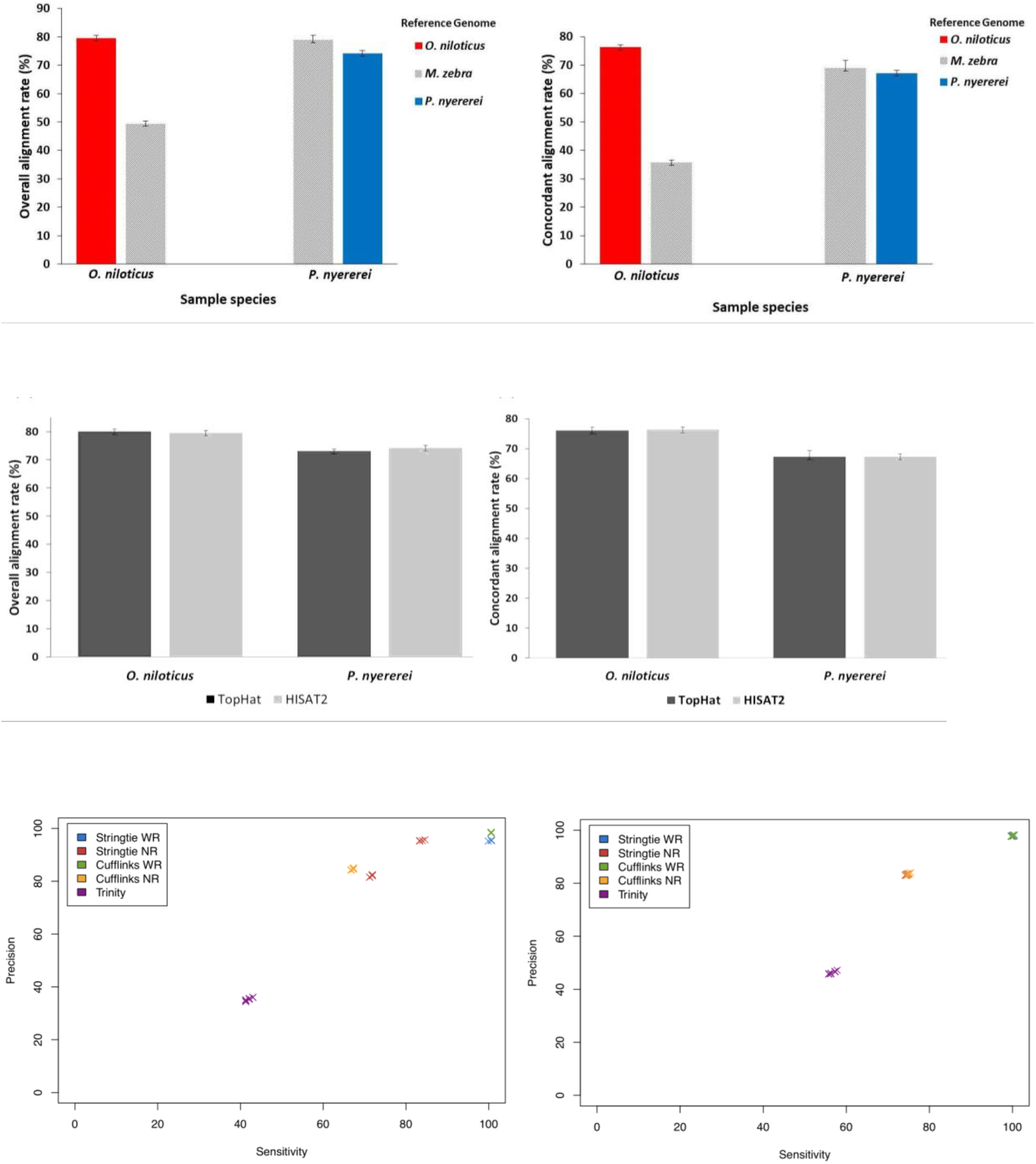
Proteogenomic workflow. 1a: Data samples procured and analysed in this study. (*Total RNA was extracted from testes of 4 PN samples) (Image of ON fish is taken from *Biolib.cz (Klas Rudloff)* and image of PN is from *african-cichlid.com*) 1b. Pictorial representation of the methods followed in the analysis. DE: differential expression. PCA: principal component analysis. GO: gene ontology. Details of individual steps given in text. 1c. Comparison of RNA-seq read alignment rates to the M. zebra, ON and PN genomes. X axis: the species from which the RNA-seq reads were derived. Colours: the genome to which the RNA-seq reads were aligned. Red: ON. Gray: M. zebra. Blue: PN. Error bars: standard errors. Figure on the left: Overall alignment rates; on the right: Concordant alignment rates. 1d. Comparison of liver RNA-seq read alignment rates using HISAT2 and TopHat. Rates of alignment of O. niloticus and P. nyererei liver RNA-seq reads to their respective genomes using HISAT2 2.1.0 and TopHat 2.1.0. Dark gray: TopHat. Light gray: HISAT2.Error Bars: standard errors. Figure on the left: Overall alignment rates; on the right: Concordant alignment rates. 1e. Sensitivity and precision of transcriptome assembly of simulated reads. Simulated reads with uniform expression levels and no sequencing errors. Sensitivity was assembled using five transcriptome assembly methods. 10×10,000 transcripts were randomly sampled with replacement from each simulated transcriptome and the sensitivity and precision of these subsets assessed using GFFcompare for O. niloticus-derived reads (figure on the left) and for the P. nyererei derived reads (figure on the right).

## Results

### Selection of the reference genome and evaluation of read alignment and transcript assembly methods

The reference genomes of ON and PN feature gaps and mis-assemblies, and are not completely annotated. This made it necessary to first examine the extent to which the poorer quality of existing assemblies for these species might affect alignment and quantitation of RNA-seq reads. We aligned the reads to their respective genomes and to the better annotated genome of a closely related cichlid species, *Metriaclima zebra*, with fewer gaps and mis-assemblies. We then compared overall and concordant alignment rates. PN liver reads had 4.9% and 1.8% higher overall and concordant alignment rate, respectively, to the *M. zebra* genome than to its own genome. Whereas, ON liver reads had 30% and 40.5% lower overall and concordant alignment rates, respectively, to the *M. zebra* genome than to its own genome **(Fig. 1c)**. As ON reads had a higher overall and concordant alignment rate on aligning to its own genome, we decided to align the reads to the species’ respective genomes for transcriptome alignment and assembly. The PN derived reads had a higher alignment rate to *M. zebra* than itself, while it was lower in ON derived reads, may be because *M. zebra* is more closely related to PN than ON. The two commonly used RNA-seq read alignment methods: TopHat and HISAT were compared by aligning the liver tissue reads of both the species to their respective genomes. The overall alignment **(Fig. 1d left)** and concordant **(Fig. 1d right)** alignment rates for both methods were very similar, but HISAT2 took approximately half the computational time compared to TopHat. Hence, HISAT2 was chosen to align the reads for the rest of the analysis.

We then evaluated several assembly methods (**Fig. 1b**). As there is no consensus in the literature regarding the optimal method for transcriptome assembly ^50 51 52 53^, the following assembly methods were evaluated: Trinity - a *de novo* method, and Stringtie and Cufflinks - two reference based methods. These two reference-based methods were run in two modes: with and without providing the reference annotations (Stringtie/Cufflinks WR and NR respectively). To compare the assembly between the methods three replicates of simulated reads were generated for both the species, using a built-in differential expression model of Polyester v1.14.1. Reads were simulated from ON and PN reference annotation transcripts to produce three replicates of approximately 25 million 75 bp paired-end simulated reads for each species, without incorporating sequencing errors and with uniform transcript expression levels. Simulated reads were aligned to their respective genomes using HISAT2 2.1.0 and then assembled using the five assembly methods. For both ON and PN, the *de novo* method, Trinity, had much lower precision and sensitivity in assembling transcripts than the reference-based methods. The reference-annotation based methods (Stringtie WR and Cufflinks WR), which used the existing genome annotations in transcriptome assembly, showed the highest precision and sensitivity values for both ON and PN. For the PN reads there was no difference between these two methods. However, Stringtie NR had higher mean precision and sensitivity values than Cufflinks NR when assembling ON-derived reads **(Fig. 1e)**. On the basis of these results, three methods were chosen for assembling the RNA-seq reads: Trinity (TR), Stringtie WR (WR) and Stringtie NR (NR). Trinity was chosen despite its low sensitivity as it was the only method studied that is capable of assembling transcripts that are not present in the reference annotations.

The Stringtie assembled transcriptomes were quantified using Stringtie to generate transcript level abundances. Whereas, RSEM was used to quantify the transcripts assembled *de novo* by Trinity. The transcripts abundances for all the three methods were then analysed for differential expression using Ballgown and Ebseq. RSEM generated counts for Trinity assembled transcripts were converted to FPKMs required for downstream assembly of Ballgown, using ‘ballgownrsem’ function.

### Identification of orthologous and uniquely expressed transcripts in the two fishes

To identify what transcripts were ‘differentially expressed’ between the two species, we had to first identify how many were common between them, for that we had to identify the orthologues as discussed below. Transcripts that are ‘uniquely’ expressed in the two species may explain the phenotypic diversity if they are functional, hence we identified ‘uniquely’ expressed transcripts between the two species as described below.

After the assembly of aligned reads with the three assembly tools; the assembled transcriptomes were processed to remove the unexpressed, duplicate and highly similar transcripts **(Supplementary Figure 1)**. Post the filter, the total number of transcripts assembled by the three assembly methods, in each tissue of each fish, as depicted in **(Fig. 2a)**, ranged from 22,879 to 254,399. The assembled transcriptomes were compared between the two fishes for each method and tissue type, to identify the transcripts that were either conserved between the two fishes or were expressed only in one or the other fish.

**Figure 2:**
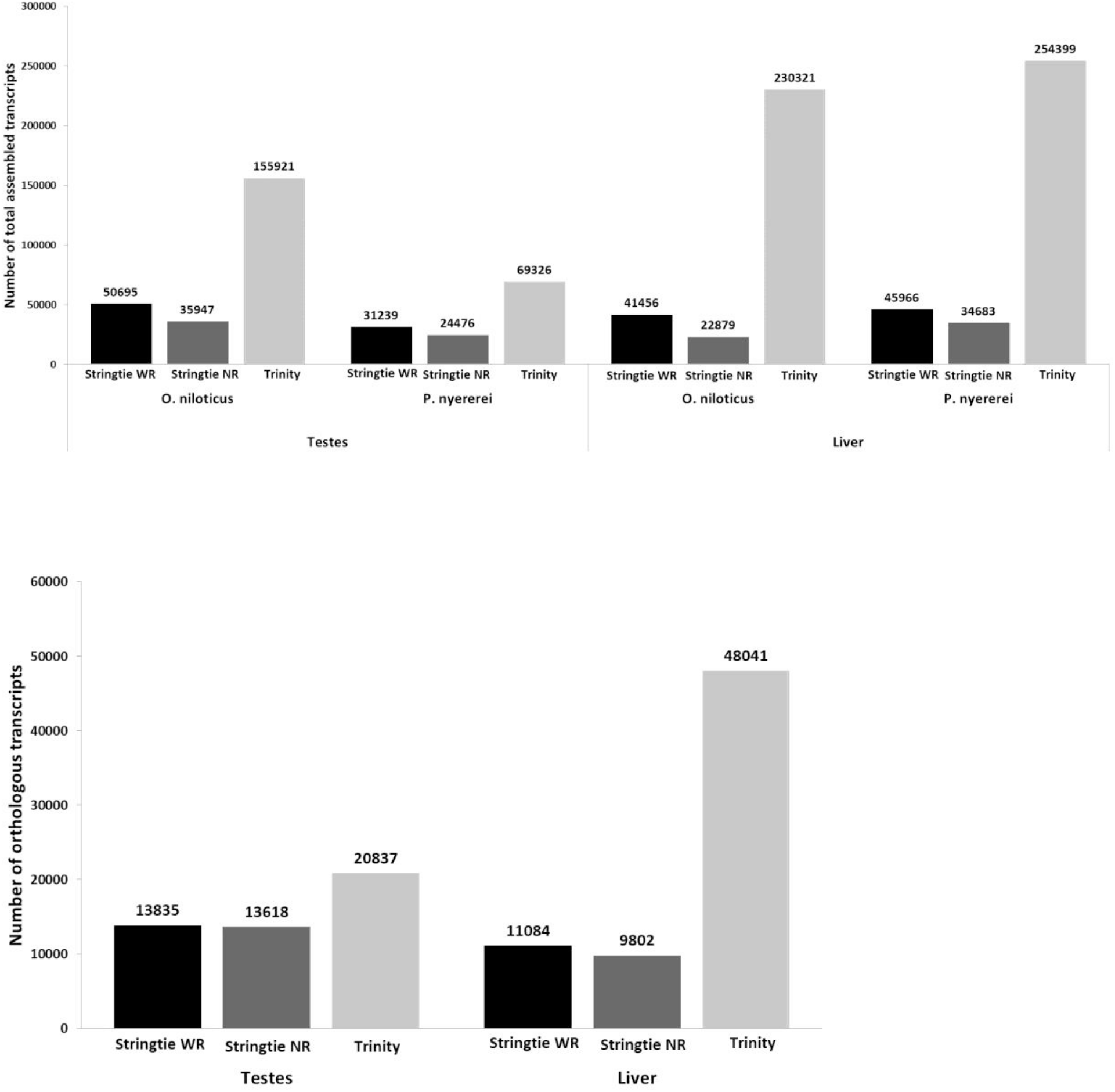
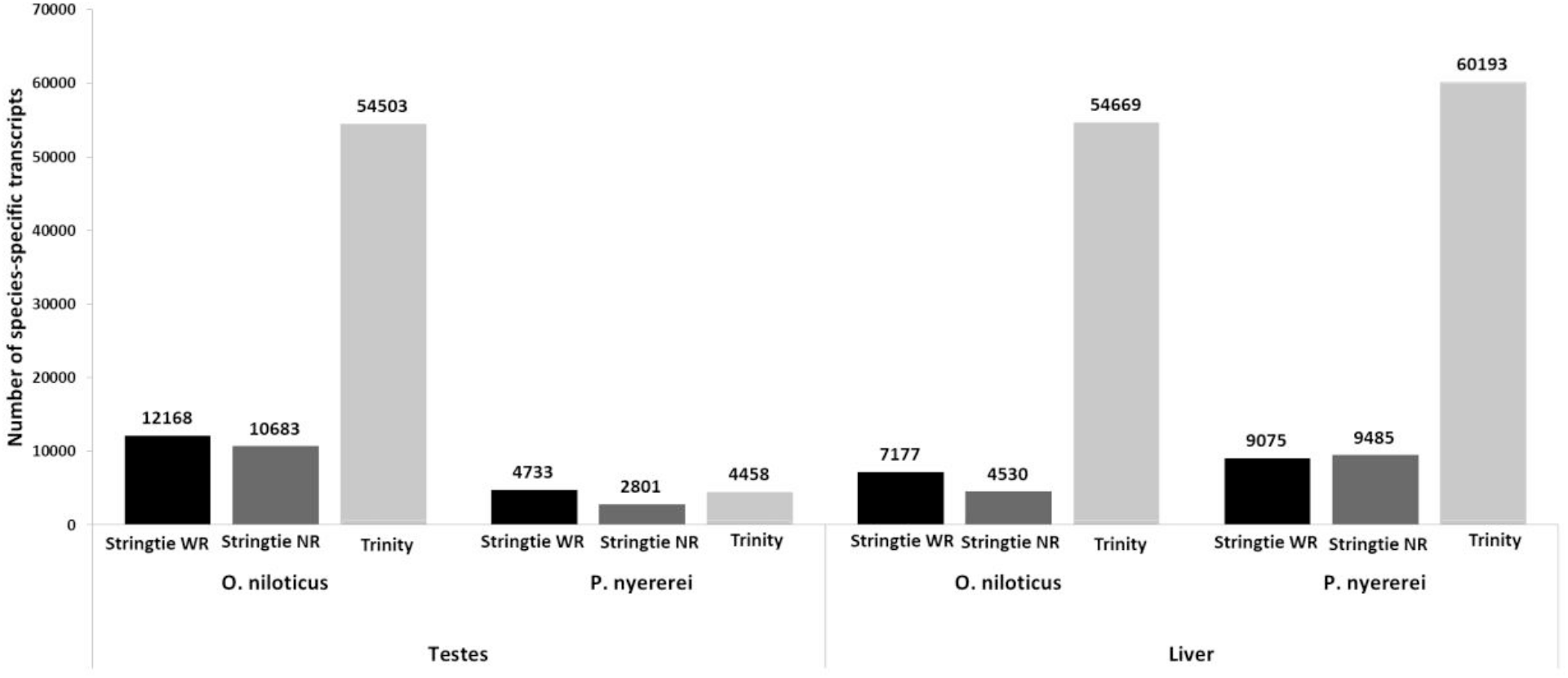
Number of transcripts identified by the three assembly methods: Stringtie WR (black), Stringtie NR (dark gray), Trinity (light gray) 2a. Total number of assembled transcripts found in each tissue of each fish for the three transcriptome assembly methods: 2b. Total number of orthologus transcripts between the two fishes in both the tissues. 2c. Number of transcripts expressed uniquely to a fish in each tissue.

We identified the ‘orthologous’ transcripts conserved between the two fishes using reciprocal best hits (RBH) method. The ON transcript sequences for each tissue type and method were mapped to their respective PN transcriptomes and vice versa using blastn v2.7.1+ ^54^. Transcript pairs that were each other’s highest scoring match were identified as orthologs. And the transcripts if did not have a match in the opposing species with at least 80% identity, were assumed to be expressed uniquely to the species; and were classes as ‘species-specific’ transcripts.

We identified, for the three assembly methods, around 13,618 - 20,837 orthologous transcripts, in the testis’s transcriptomes of the two fishes. Similarly, around 9,802 - 48,041 orthologous transcripts were identified in the liver transcriptomes of the two fishes **(Fig. 2b)**. In both tissues, the number of orthologous transcripts identified in the Trinity assembled transcriptomes were highest, while were lowest in the Stringtie NR assembled transcriptomes.

Additionally, **Fig. 2c** depicts the number of species-specific transcripts identified by three assembly methods, per tissue in each fish. The number of ON-specific transcripts, in the two tissues for the three methods, varied from 4,530 - 54,669, whereas PN-specific transcripts varied from 2,801 - 60,193. Except for PN testes, the number of species-specific transcripts identified in Trinity-assembled transcriptome was highest. No species-specific liver transcripts were commonly found by all three of Stringtie WR, Stringtie NR and Trinity. In contrast, 441 (0.6%) of the species specific ON testes transcripts and 93 (0.9%) of the species-specific PN testes transcripts were identified by all three methods **(Supplementary Figure 2)**.

### Comparative transcriptomes between liver and testes of the two fishes

To determine whether the transcriptome level differences contribute to the diversity in the two fishes, we compared the expression levels of the orthologous transcripts of the equivalent tissues. Principal component analysis (PCA) on the normalised expression levels qualitatively separated the two fishes in both the liver and testes samples for all three transcriptome assembly methods **(Figure 3)**. Next, we carried out differential expression analysis of the orthologous transcripts to identify transcripts whose expression varied between the two fishes using the both Ballgown and EBseq tools. Ballgown analysis revealed that no transcripts were differentially expressed between the liver and testes transcriptome of the two fishes. But, when differential expression analysis was done using Ebseq, transcripts from both testes and liver were identified to be DE. 4,591-26,671 and 8,872-13,436 transcripts were identified to be respectively DE in liver and testes transcriptomes assembled by the three assembly pipelines. As large numbers of orthologous transcripts (~30-62%) were identified to be DE by EBseq, we did not further analyse these results.

**Figure 3.**
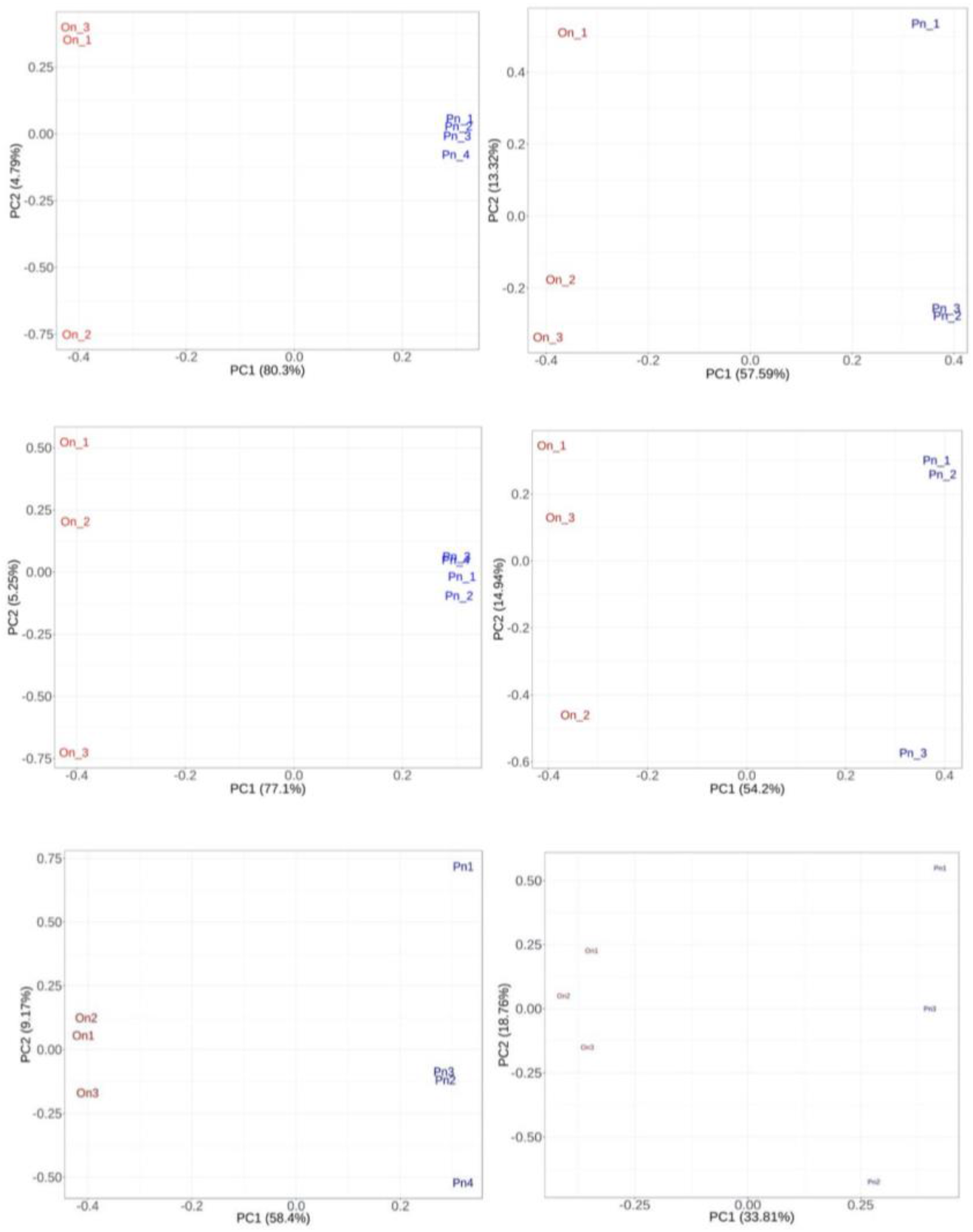
Principal component analysis plots for each tissue type and transcriptome assembly method of the samples separated based on FPKM values of orthologous transcripts. (A) Stringtie WR testes. (B) Stringtie NR testes. (C) Trinity testes. (D) Stringtie WR liver. (E) Stringtie NR liver. (F) Trinity liver. In both the tissues, for all the three assembly methods, the samples for two fishes separate over the first principal component and form separate clusters.

### Functional annotation of the species-specific transcripts

For annotation by Interproscan and blastp we used union of the transcripts identified by the three assembly methods, and not just the transcripts identified commonly by the three methods. Functional annotations of species-specific transcripts, for each tissue type and species showed broadly similar trends, mostly pertaining to cellular processes, metabolic processes, localisation and regulation of biological processes. Some annotations were also specific to a particular tissue type. Eight of the species-specific transcripts in the PN testes and fifteen of the species-specific transcripts in the ON testes were annotated with the GO term reproductive processes and reproduction, suggesting that the ON and PN reproductive systems have diverged (**Supplementary Figure 3**).

### Identification of the novel transcripts derived from the noncoding regions

Further analysing the species-specific transcripts, we observed that a subset of them were transcribed from previously unannotated noncoding regions. We call these regions novel Open Reading Frames (nORFs). Of these subset, 100 nORF transcripts had evidence of translation identified using our mass spectrometry-based proteogenomic analysis. We observed that 8-24 and 5-25 number of nORF transcripts were transcribed and translated from intronic and intergenic regions respectively, found for each species and tissue type **(Table 1)**. There was little overlap in the species-specific nORF translated products found by each method, with no overlap between Trinity and the Stringtie methods, two intronic and two intergenic species-specific liver ON translation products found by both Stringtie WR and Stringtie NR, six and two intergenic species-specific liver PN and testes ON translation products found using both Stringtie WR and Stringtie NR.

**Table 1.** Identification of novel ORFs. The number of unique intergenic and intronic novel species-specific translation products for each tissue and species

|  | Intergenic | Intronic |
| --- | --- | --- |
| O. niloticus testes | 12 | 24 |
| O. niloticus liver | 12 | 14 |
| P. nyererei testes | 5 | 0 |
| P. nyererei liver | 25 | 8 |
|  | **54** | **45** |

Further investigation of these nORF translated products by InterProScan revealed that one intergenic product from PN testes was annotated with immunity related GO terms. Similarly, one intergenic translated product, each from ON testes and PN liver, had immunoglobulin like fold and domain.

### Evolutionary analysis of nORF transcripts

In order to determine whether these 100 nORF transcripts, with direct evidence of translational evidence because of the presence of peptides - detected by mass-spectrometry analysis, evolved in a non-neutral manner we next calculated their substitution rates by calculating the genome-wide, base-wise conservation-acceleration (CONACC) scores using phyloP ^34^. To do this, existing multiple whole genome alignments of the five cichlids provided by Brawand et al ^33^ was used. Of the 100 nORF transcript regions; we were able to map the scores for only 41 regions because of the variability in the two different ON assemblies and due to insufficient aligned data. As the ON assembly, ASM185804v2, used in our analysis was different than the one used in the whole genome alignments - Orenil1.1, few of the nORF regions were unmapped during assembly conversion. The regions with the mapped scores were further reduced as no CONACC score is assigned to a site; if there is insufficient data per site or gaps in the alignment **(Table 2)**.

**Table 2:** Number of novel transcripts that were mapped between the ON’s two assembly versions and later with CONACC scores.

|  | Identified in our study<br>(ASM185804v2 /<br>PunNye1.0) | On mapping to newer<br>assembly (OreNil2 /<br>PunNye1.0) | Mapped with<br>CONACC scores |
| --- | --- | --- | --- |
| ON novel intergenic | 24 | 15 | 9 |
| ON novel intronic | 38 | 35 | 27 |
| PN novel intergenic | 30 | 30 | 0 |
| PN novel intronic | 8 | 8 | 5 |

CONACC scores were computed over all branches of the cichlid’s phylogeny, and used to detect the departure from neutrality in novel regions and also in the other known annotated features of the genome like CDS, 5’UTR, 3’UTR, introns, intergenes and ancient repeats (AR). The analysis of the cumulative distributions **(Fig. 4a)** of the phylop scores of ON’s known annotated features showed that the CDS regions (red line) were most conserved while the AR’s were least conserved. This is intuitive as the functional coding regions are expected to have more evolutionary constraints than the non-functional repeat regions. The distribution of CONACC scores of all the annotated features were significantly different than that of AR (Welch t-test, P-value < 0.05) **(Fig. 4a)**.

**Figure 4:**
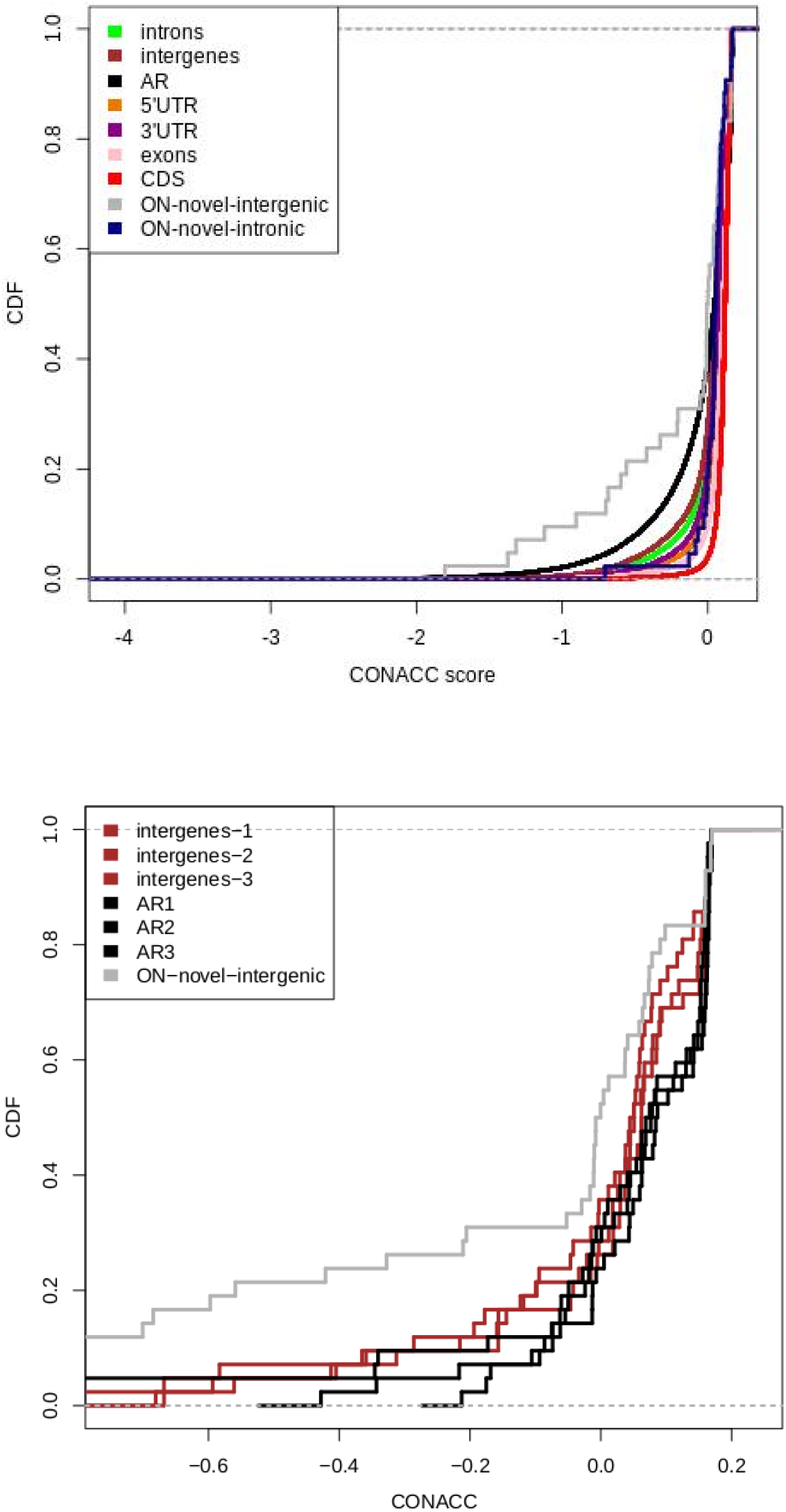
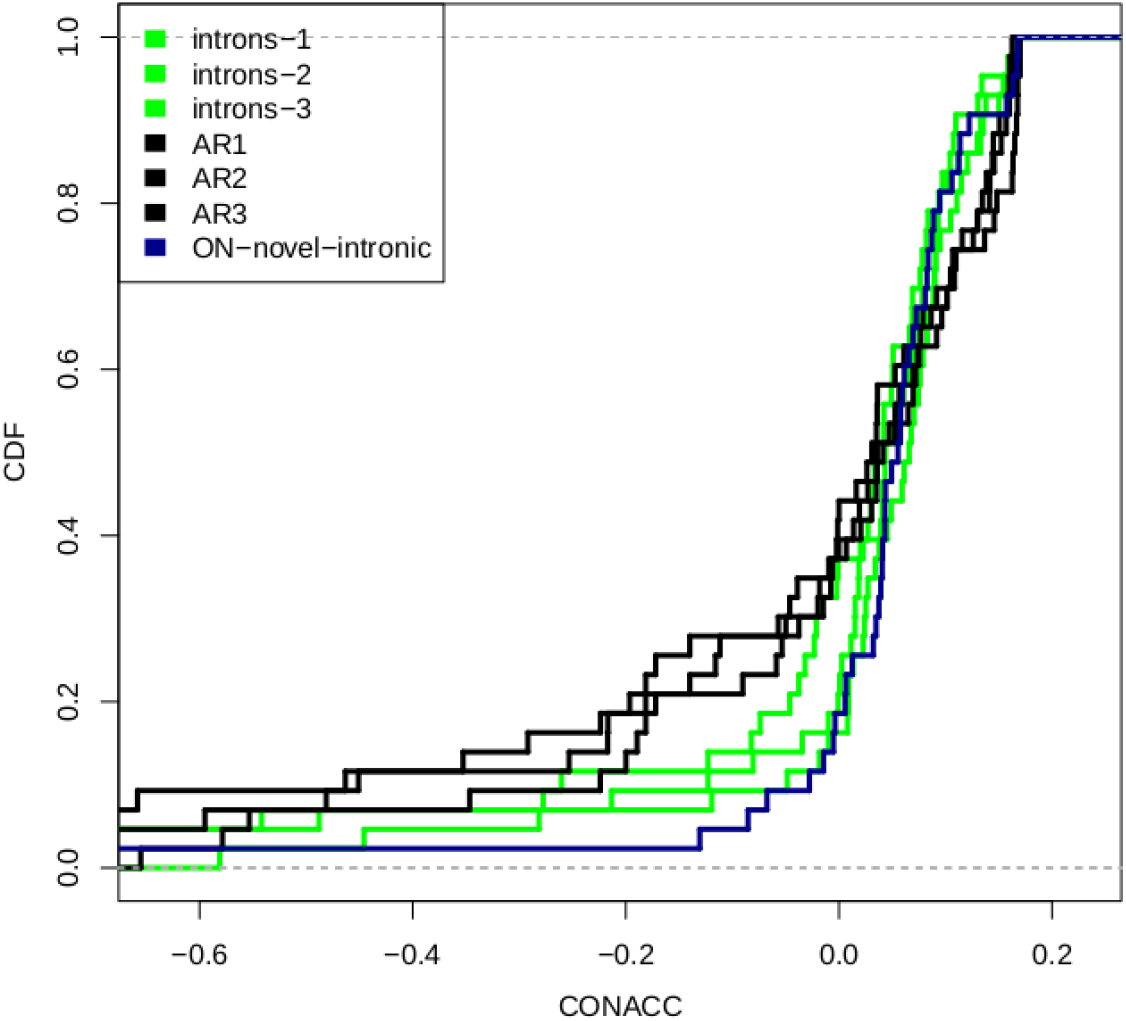
Distribution of conservation-acceleration (CONACC) scores calculated using phyloP over all-branch analysis including 5 cichlids for: (A) Different features of ON’s genome. AR - ancestral repeats, 5’UTR - 5’ untranslated region, 3’UTR - 3’ untranslated region, CDS - protein coding sequences. The distribution of CONACC scores for all the features is significantly different than that of AR (Welch t-test, P-value < 0.05) (B) Three sets respectively of randomly-picked, AR regions (black) and intergenic regions (brown), which are length-matched and are equal sample-sized to the novel intergenic regions. The distribution of CONACC scores of the randomized AR subsets were significantly different from that of novel intergenic regions for 7519/10000 times. (C) Three sets respectively of randomly-picked, AR regions (black) and intronic regions (light green), which are length-matched and are equal sample-sized to the novel intronic regions. The distribution of CONACC scores of the randomized AR subsets were significantly different from that of novel intergenic regions for 2338/10000 times.

Conservation scores were also mapped to the 9 ON novel intergenic and 27 ON’s novel intronic regions. As these novel regions are very few compared to the AR, we sampled 10,000 times, from all the AR regions, to randomly pick one length-matched AR per nORF transcript. The distribution of CONACC scores for these length-matched, equal sample-sized AR regions were significantly different (Welch t-test, p-value < 0.05) than the novel intergenic regions (**Fig. 4b**) for 7,519/10,000 times; and only 2,338/10,000 times for the novel intronic regions **(Fig. 4c)**.

Compared to AR, the 9 novel-intergenic regions in ON showed a shift towards more accelerated CONACC scores (gray line in the graph), whereas the 27 novel-intronic regions showed a non-neutral substitution rate with shift towards more conserved CONACC scores (blue line in the graph). This indicates that these regions which are varied in all the cichlids, might contribute to the phenotypic variation in ON.

### Phylogenetic divergence time scale analysis of ON and PN

To check whether these accelerated nORF genomic regions can reveal the actual divergence time between ON and PN species and perhaps give us a clue to the speciation process we carried out Bayesian Evolutionary Analysis Sampling Trees model (BEAST) ^35^, which was run on BEAST v1.10.4 ^55^. A strict molecular clock was set, to allow for the most reliable comparison between trees based of nORFs sequences. The molecular clock was time calibrated with a fossil time constraint. The constraint was set as a lognormal prior distribution with a mean in real space of 45.5 million years ago (MYA) and a standard deviation of 0.5 MYA for all the entire group of cichlids. This time calibration was based on cichlid fossils estimated to be 45 million years old ^56^. The fossil was recovered from Mahenge in the Singida region of Tanzania. The reason why we chose this fossil estimate is because according to Murray, 2001 ^56^ not only are the Mahenge cichlids the oldest known species but, as a potential flock, they are the oldest record of any kind of species flock formation in the Cichlidae. Other fossil cichlids from an Oligocene lake in Saudi Arabia were considered as belonging to several different lineages and, therefore, do not constitute a species flock. The Mahenge cichlids, therefore, provide the first fossil evidence to indicate that the ability of the cichlids to form species flocks arose prior to 40 Myr ago. The substitution rate was fixed to allow better comparison between trees.

We carried out ten phylogenetic trees analysis based on bayesian inference to assess whether these nine nORFs showed recent divergence (**Table 3 and Figure 5**). In addition to the nine nORFs we used two control genes that would have arisen before the divergence. They were, the DNA methyltransferase 1 gene (DNMT1) which encodes for DNA (cytosine-5)-methyltransferase 1 enzyme, which is essential for DNA cytosine methylation and therefore would act as a housekeeping gene, and an ancestral repeat as they are highly conserved between the different fish species and would be expected to have a low mutation level.

**Table 3:** The relative divergence time between O. niloticus and P. nyererei, along with the deviation from the Neutral Model that was calculated using CONACC Scores, which tissues these nORF’s where found in, what pipeline was used to assemble the transcripts and whether these nORFs shared any domains with known proteins. These nORFs were identified in the intergenic region of *O. niloticus*.

| Sequence | Relative Divergence Time (MYA) | CONACC Score | Tissue | Transcript Assembly Pipeline | InterPro Scan |
| --- | --- | --- | --- | --- | --- |
| DNMT1 | 52.22 | 0.0000 | NA | NA | NA |
| (AR) LG18:20803645-20804072 | 104.58 | 0.1055 | NA | NA | NA |
| LG5:36477108-36477531 | 154.82 | 0.0638 | Liver | Trinity | - |
| LG2:4126626-4126944 | 160.26 | 0.0368 | Testes | Trinity | - |
| LG6:20251315-20251636 | 233.87 | -0.0302 | Testes | Trinity | - |
| LG17:13477145-13481614 | 37.98 | -0.3523 | Testes | Trinity | - |
| LG5:21555805-21556303 | 31.07 | 0.0862 | Testes | Stringtie NR | - |
| LG15:24005494-24006333 | 9.6 | 0.0284 | Liver | Trinity | - |
| UNK275:21277-21517 | 2.97 | -0.0003 | Liver | Stringtie NR | - |
| UNK219:32598-35307 | 182.04 | 0.0845 | Testes | Stringtie NR | FGFR1 ONCOGENE PARTNER/LISH DOMAIN-CONTAINING PROTEIN |

**Figure 5.**
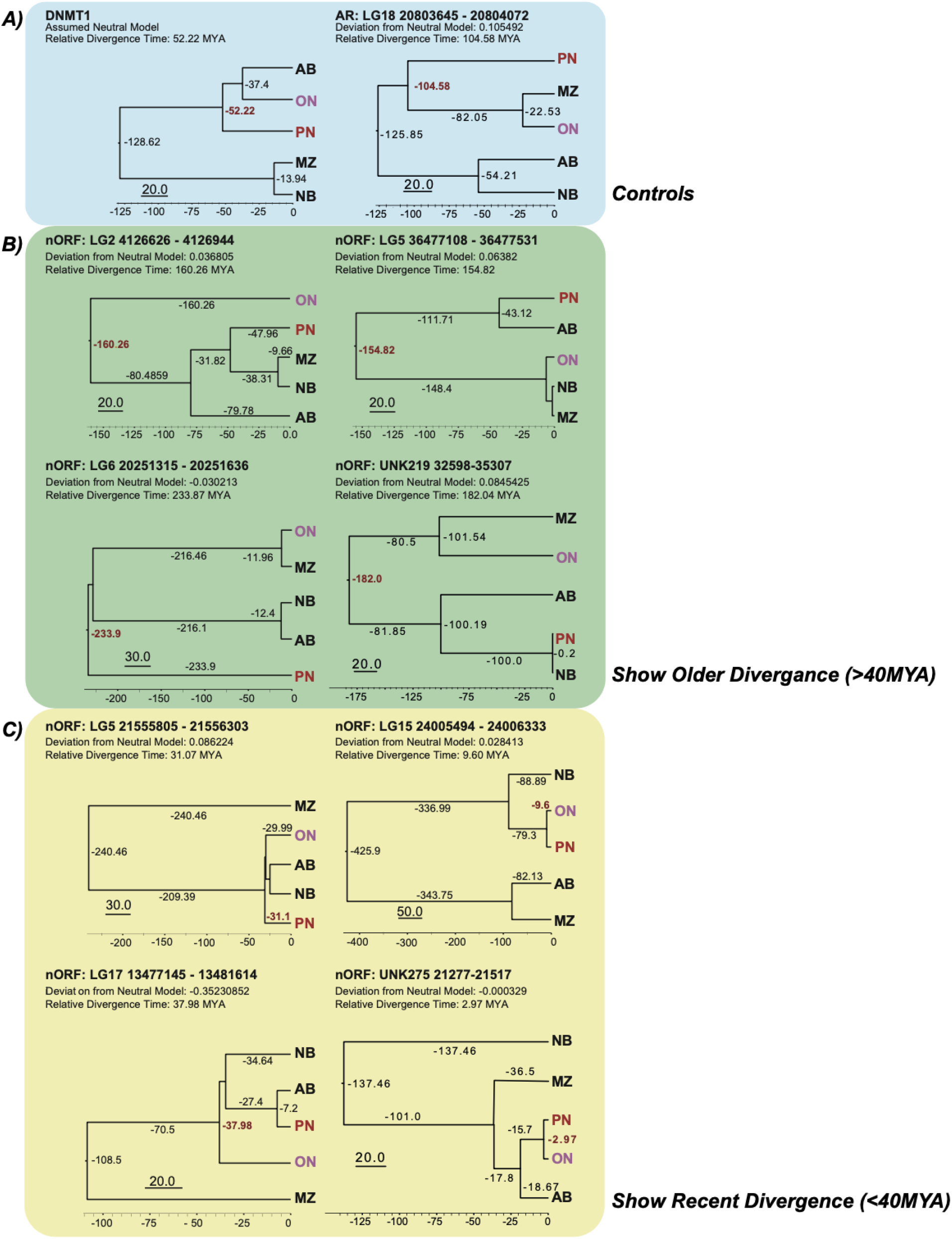
Phylogenetic trees based on nORF sequences that were time-calibrated using fossil priors and substitution rate. These trees were constructed using BEAST v1.10.4. The DNMT1 gene and AR (ancestral repeat) were used as controls. The nORF’s selected had been shown to deviate from the Neutral Model. For each tree we show the degree of deviation from the Neutral Model, along with the relative divergence time between PNi and ON, that was calculated based on that particular sequence and deviation from the Neutral Model (which was set at 1). A) DNMT1 and ancestral repeat sequence divergence used as controls. B) Four nORFs that showed divergence prior to 40 MYA. C) Four nORFs that showed divergence earlier than 40 MYA.

The coordinates of these nine nORFs were entered into the Cambridge Cichlid Genome browser using ON (Broad OreNil1.1/OreNil2 assembly) as the reference point. This tool was used to extract the orthologous sequences for the other four cichlid species including PN, using the 8-way comparative genomic track setting option. The sequences for the DNMT1 gene were all extracted manually from the National Center for Biotechnology Information (NCBI) database for each species. The relative divergence time of the nORFs and the controls between the two cichlid fishes ON and PN was calculated based on bayesian inference analysis.

The phylogenetic trees were constructed using five cichlid species with genomes that were published in the Brawand et al., 2014 ^33^ and we just focus on the divergence of the selected genome coordinates (as discussed in the methods section).The phylogenetic trees showed 52.22 and 104.58 MYA divergence for DNMT1 and the ancestral repeat respectively (**Figure 5A**), which is greater than the Mahange cichlids fossil’s age. Two of the nine nORFs were actually the same nORF, which is found in the UNK219 32598-35307 region and therefore only one phylogenetic tree was constructed for this nORF, as it was identified twice. The four nORF phylogenetic trees shown in **Figure 5B**, show divergence greater than 40 million years, which is more than the age of the Mahenge fossils; therefore, these nORFs are likely to not contribute to the speciation of cichlids. **Figure 5C**, shows the remaining four nORFs that exhibit a recent relative divergence time. These four nORFs show relative divergence times between ON and PN that vary from 3 to 38 million years ago. We postulate that these nORFs might play a role in the adaptation of these cichlids which allows them to undergo adaptive radiation.

## Discussion

Despite the differences caused by different transcriptome assembly methods, some biological results were found independent of the method used. Transcript and protein expression levels had not diverged in the liver and testes, despite known differences in the diets of ON and PN, which we expected to affect liver gene expression. However, the presence of species-specific transcripts in testes indicate that the tests’ transcriptome is undergoing rapid change during the evolutionary process. Eight of the species-specific transcripts in the PN testes and fifteen of the species-specific transcripts in the ON testes were annotated with the GO term reproductive processes and reproduction, suggesting that the ON and PN reproductive systems have diverged (**Supplementary Figure 3**).

There are such similar observations made for other taxa too. For example, Jagadeeshan et al., 2005 ^57^ found more non-synonymous substitution in testes-expressed genes of Drosophila than in genes expressed in the ovaries and head tissues. Similarly, Voolstra et al, 2007 ^58^ found greater expression divergence in the testes compared to the brain, liver and kidneys when comparing mouse species and Khaitovich et al., 2005 ^59^ had similar findings with regards to humans and chimpanzees. Jagadeeshan et al., 2005 ^57^ hypothesised that the greater divergence in sex traits than non-sex traits in Drosophila may relate to the establishment of reproductive isolation (RI) during speciation. This is, however, unlikely to be the case in cichlids as there is little post-zygotic RI in closely rated cichlid species, with pre-mating RI predominating ^60^. The divergence in testes gene expression could alternatively relate to sperm competition between males, a phenomenon which is common in polygynous species with maternal care of the young and which is intensified by mouth brooding, a trait which is common to both species ^61,62^. Differences in the response to sperm competition in the two species could account for the differences in testes gene expression. Some of the differences in testes gene expression could also relate to changes in sex determination systems. Sex determination is a labile trait within the cichlids and the downstream mechanisms for sex determination are less conserved in the cichlids than in other taxa ^63^. As the downstream pathways are expressed in the gonads this may contribute to the species-specific expression of transcripts in testes’ of ON and PN. Indeed the question of the interaction and of the relative importance of natural and sexual selection in the adaptive radiations of cichlid fishes remains unanswered ^39^. However, Wagner et al ^1^ claim that extrinsic environmental factors related to ecological opportunity and intrinsic lineage-specific traits related to sexual selection both strongly influence whether cichlids radiate.

The proteogenomic analysis that we performed by integrating the transcriptomic and proteomic data from liver and testes tissues of ON and PN demonstrated the existence and expression of nORFs from the intergenic and intronic regions that were not previously observed. Further investigation of these nORF translated products by InterProScan revealed that one intergenic product from PN testes was annotated with immunity related GO terms. Similarly, one intergenic translated product, each from ON testes and PN liver, had immunoglobulin like fold and domain. Because it was not possible to assess the functional role of these newly diverged nORF regions, we performed evolutionary and phylogenetic analysis of these nORFs (**Table 2 and 3 and Figure 4 and 5**) with statistical validation, which revealed that some nORFs emerge from the ‘accelerated’ regions of the genomes. More importantly, four of the eight nORFs that emerged from the ‘accelerated’ regions of ON indicated a recent divergence time of 3-38 million years from PN. Brawand et al ^33^ assessed that ON and PN diverged approximately in the same time scale. We believe that the similarity in time scales may not be a fortuitous coincidence.

Our results indicate that it is possible to partially explain the rapid speciation of cichlids fishes in general if we systematically explore, identify and analyse nORF regions in every species. As a limitation of this study we are indeed aware that phylogenetic trees for groups of closely related species often have different topologies, depending on the genes (or genomic regions) used ^64^. The translated products from these nORFs may not have yet evolved functions leading to their fixation. Perhaps, they are still being ‘tinkered’ with the potential to optimize, or perhaps even change. The emergence of these nORFs is intriguing and we postulate that they might evolve into functional genes contributing to the speciation of the cichlids fishes. Therefore, this study supports the hypothesis that de novo emergence may be the dominant mechanism of novel gene emergence and perhaps they may contribute to increased fitness, as they can become essential. This study also suggests that population biodiversity can be brought about by rapid evolution of intergenic genomic regions.

Organisms that undergo extensive speciation with diverse phenotypic variation, such as the cichlids, must have highly ‘evolvable’ genomes - especially genomes that are more evolvable in the noncoding regions than in the coding regions because all the known proteins eventually tend to get fixed over geological time-scales. We have presented evidence that shows the presence of evolutionary-accelerated regions in certain noncoding regions that exhibit coding potential and suggest that this may be a potential cause of speciation. One study has demonstrated that some noncanonical proteins evolve neutrally, and that can at some point they acquire new functions ^28^. Another study has demonstrated that nORFs pervasively emerge from noncoding regions but are rapidly lost again, while only a relatively few are retained much longer ^29^. We believe we have answered the central question that we set out to answer - whether nORFs can emerge from intergenic regions. Our study also provides evidence for the correlation of nORF emergence and speciation and divergence of ON and PN, systematic investigation of which may reveal more clues in future studies. But the most important question as to whether transcription emerged first or whether nORFs emerged first from intergenic regions remains to be answered.

## Data availability

The mass spectrometry proteomics data have been deposited to the ProteomeXchange Consortium via the PRIDE partner repository with the dataset identifier PXD019072. RNAseq data can be downloaded from GEO: GSE150744 https://www.ncbi.nlm.nih.gov/geo/query/acc.cgi?acc=GSE150744

## Acknowledgements

We would like to thank Seven Bridge Genomics (https://www.sevenbridges.com/) for letting us use their cloud platform. We would like to thank RosettaHuB (https://rosettahub.com) for helping us to build applications using Amazon Web Services. We would like to thank Dr. Welch for help with the BEAST analysis. The authors would like to thank Ole Seehausen and Marcel Häsler (Universität Berne) for providing *P. nyererei* tissues, David Penman (University of Stirling) for providing *O. niloticus* tissues, Mingliu Du for help with RNA extractions and the sequencing facility at the Wellcome Sanger Institute for NGS libraries preparation and RNA sequencing. We thank the anonymous reviewers and editor for fantastic critical comments that improved the manuscript greatly.

## Author contributions

Sh. P. did the transcriptomic, evolutionary and over all data analysis and gave inputs for writing the manuscript. RN did the initial transcriptomic and proteogenomic analysis. JSMN did the time-calibration analysis. RC did the proteomics experiments. TW did the initial quantitative transcriptomic analysis. MTW did the initial transcriptomic analysis. YU participated in the proteogenomics analysis. GV and EAM generated the ON and PN liver transcriptomic datasets. SP designed and supervised the project, analysed the data, and wrote the manuscript.

## Materials and Methods

### Fish samples

*P. nyererei* (generation 1; Lake Victoria) and *O. niloticus* (generation ~93; Manzala, Egypt) liver and testes samples were obtained in accordance with the relevant guidelines and regulations of the University of Cambridge by the Miska lab. This RNA and protein extraction methodology was approved by the University of Cambridge.

### Total RNA extraction from liver tissues and sequencing

Approximately 5-10mg of fresh liver tissue from tank-reared *P. nyererei* (generation 1; Lake Victoria) and *O. niloticus* (generation ~93; Manzala, Egypt) specimens, snap-frozen upon dissection, was homogenised and used for RNA extraction. Total RNA was isolated using TRIzol (ThermoFisher) and then treated with DNase (TURBO DNase, ThermoFisher) to remove any DNA contamination. Quality and quantity of total RNA extracts were determined using NanoDrop spectrophotometer (ThermoFisher), Qubit (ThermoFisher) and BioAnalyser (Agilent). Following ribosomal RNA depletion (RiboZero, Illumina), stranded rRNA-depleted RNA libraries (Illumina) were prepped and sequenced (paired-end 75bp-long reads) on HiSeq2500 V4 (Illumina) by and at the Sanger Sequencing Facility. On average, 11.82±0.42Mio paired-end reads were generated for ON and PN liver samples.

### Extraction of ON and PN total cell proteome

Liver samples from the same ON and PN fishes that were used for RNA extraction were used for the mass-spectrometry analysis as well. A new set of ON and PN fishes were used to obtain testes samples. To extract the total cellular proteome, ~5 mg of tissue were lysed in buffer (6M Urea, 2M Thiourea, 4% CHAPS, 5mM Magnesium Acetate, 30mM Tris pH 8.0), and 15μg protein in 5x Laemmli buffer with 5% b-mercaptoethanol was loaded on Mini-PROTEAN TGX Precast Gels (BioRad). Gel lanes were cut into three sections for peptide extraction. Gel sections were cut into 1-2mm cubes, washed with 50% Acetonitrile and 100mM Ammonium bicarbonate solution until blue stain is washed. Gel pieces were treated with 100% Acetonitrile, and then reduced with 10mM DTT in 100mM Ammonium bicarbonate for reduction at 56°C for 1 hour, and alkylated with 55mM Iodoacetamide in 100mM Ammonium bicarbonate in dark for 45 min at room temperature. Gel pieces were washed with 100mM Ammonium bicarbonate, and then treated with 50% Acetonitrile followed by 100% Acetonitrile. Subsequently, gel pieces were treated with diluted trypsin (5ng/ul) enzyme, overnight at 37°C. Peptides were extracted, dried, and dissolved in 3% Acetonitrile with 0.1% Formic Acid. A total of 36 total samples (2 fishes * 2 tissues * 3 biological replicates * 3 bands = 36) were analyzed by mass-spectrometry.

### Mass spectrometry analysis of the cichlids proteome

All LC-MS/MS experiments were performed using a Dionex Ultimate 3000 RSLC nanoUPLC (Thermo Fisher Scientific Inc, Waltham, MA, USA) system and a Q Exactive Orbitrap mass spectrometer (Thermo Fisher Scientific Inc, Waltham, MA, USA). Separation of peptides was performed by reverse-phase chromatography at a flow rate of 300 nL/min and a Thermo Scientific reverse-phase nano Easy-spray column (Thermo Scientific PepMap C18, 2microm particle size, 100A pore size, 75microm i.d. x 50cm length). Peptides were loaded onto a pre-column (Thermo Scientific PepMap 100 C18, 5microm particle size, 100A pore size, 300microm i.d. x 5mm length) from the Ultimate 3000 autosampler with 0.1% formic acid for 3 minutes at a flow rate of 10 microL/min. After this period, the column valve was switched to allow elution of peptides from the pre-column onto the analytical column. Solvent A was water + 0.1% formic acid and solvent B was 80% acetonitrile, 20% water + 0.1% formic acid. The linear gradient employed was 2-40% B in 30 minutes.

The LC elutant was sprayed into the mass spectrometer by means of an Easy-Spray source (Thermo Fisher Scientific Inc.). All m/z values of eluting ions were measured in an Orbitrap mass analyzer, set at a resolution of 70000 and was scanned between m/z 380-1500. Data dependent scans (Top 20) were employed to automatically isolate and generate fragment ions by higher energy collisional dissociation (HCD, NCE:25%) in the HCD collision cell and measurement of the resulting fragment ions was performed in the Orbitrap analyser, set at a resolution of 17500. Singly charged ions and ions with unassigned charge states were excluded from being selected for MS/MS and a dynamic exclusion window of 20 seconds was employed.

### Proteogenomic workflow to investigate evidence of translation

The 36 Thermo mass spectrometry raw files were submitted to be searched against the respective per-fish, tissue-assembled transcriptome databases (for example liver Stringtie WR-assembled, liver Stringtie NR-assembled, liver *de novo* Trinity-assembled) in six frames utilizing Proteome Discoverer v2.1 and Mascot 2.6. The spectra identification was performed with the following parameters: MS/MS mass tolerance was set to 0.8 Da, and the peptide mass tolerance set to 10ppm. The enzyme specificity was set to trypsin, and two missed cleavages were tolerated. Carbamidomethylation of cysteine was set as a fixed modification, whilst variable modifications consisted of: oxidation of methionine, phosphorylation of serine, threonine and tyrosine, and deamidation of asparagine and glutamine. High confidence peptide identifications were determined using Percolator node, where false discovery rate estimation (FDR) < 0.01 was used. A minimum of two high confidence peptides per protein was required for identification

### RNA-Seq Simulation Experiment

An RNA-sequencing experiment was simulated to assess the precision and sensitivity of *de novo* and reference-based transcriptome assembly methods ^65^ and therefore to decide which methods to use for transcriptome assembly. Reads were simulated from the *O. niloticus* and *P. nyererei* reference annotation transcripts using Polyester v1.14.1 to produce three replicates of approximately 25 million 75 bp paired-end simulated reads for each species. The reads were simulated without sequencing errors and with uniform transcript expression levels.

### Comparison of Methods for RNA-seq Read Alignment

Total RNA-seq read sequences from *O. niloticus* and *P. nyererei* testes tissues were obtained from Brawand et al. ^33^ and total RNA-seq read sequences for liver tissues were generated for this study (see above). These reads were quality-checked using FastQC v0.11.5. It was thought that the RNA-seq reads might align better to the genome of M. zebra, a closely related species, than to the *O. niloticus* and *P. nyererei* genomes, as the M. zebra genome has few gaps and mis-assemblies ^66^. The alignment rates of the RNA-seq reads to the M. zebra genome and to the species’ respective genomes were therefore compared. For this step, mapping was performed using HISAT2 2.1.0. The alignment rates of the RNAseq reads to their respective genomes using two different alignment methods: HISAT2 2.1.0 and TopHat 2.1.0 ^67^ was also compared to assess which method should be used for alignment. The read sequences from *O. niloticus* and *P. nyererei* were mapped to the reference assemblies ASM185804v2 and PunNye1.0, respectively.

### Comparison of methods for transcriptome assembly

Simulated reads were aligned to their respective genomes using HISAT2 2.1.0 ^68^ and were assembled using four reference-based assembly methods and Trinity v2.0.6 ^69^, a de novo transcriptome assembly method. The four reference-based methods were Stringtie v1.3.3 with the reference annotation (Stringtie WR), Stringtie v1.3.3 without the reference annotation (Stringtie NR) ^68^, Cufflinks 2.2.1 with reference-annotation based-transcriptome assembly (Cufflinks WR) ^70^ and Cufflinks 2.2.1 without reference annotation based-transcriptome assembly (Cufflinks NR) ^71^. The Trinity-assembled transcriptomes were mapped to their respective genomes using GMAP version 2017-11-15 ^72^ to provide genomic coordinates of the transcripts for comparison to the reference Annotations.

The precision and sensitivity of the simulated transcriptomes produced using different transcriptome assembly methods were assessed by comparison to the reference annotations. 10 × 10,000 transcripts were randomly sampled with replacement from each simulated transcriptome. These were mapped against the reference transcriptomes from which the reads were derived using GFFcompare v0.10.1 to obtain estimates for the precision and sensitivity of each assembly method at the transcript level. Raw sensitivity values were multiplied by transcriptome size / 1000 to account for the loss of sensitivity produced by using a subset of the data.

### RNA-Seq Read Alignment and Assembly

Based on the results of the simulation study, the RNA-Seq reads were assembled using Stringtie WR, Stringtie NR and Trinity. The HISAT2-aligned reads were assembled using Stringtie WR and Stringtie NR. For the reference-based assembly methods, the transcriptomes assembled for each biological replicate were merged using the Stringtie merge utility to produce one transcriptome per method per tissue per species. The Trinity assembled transcriptomes were mapped to their respective genomes using GMAP version 2017-11-15 to provide the genomic coordinates of the transcripts. This was required in order to compare the transcripts found using different methods.

### Transcriptome Processing and Database Production

The assembled transcriptomes were processed prior to data analysis and transcriptome database production. Unexpressed transcripts derived from the reference annotations were present in the Stringtie WR transcriptomes. The Stringtie-assembled transcriptomes were therefore filtered to remove unexpressed and duplicated transcripts. At some loci, Stringtie produced a large number of very similar transcripts. To reduce the number of highly similar transcripts in Stringtie-assembled loci, the Stringtie transcripts were k-means clustered within each locus and transcripts within each cluster were merged. Clustering was performed using Ballgown v2.10.0 ^68^ and transcripts were merged by taking the union of the exon coordinates of the individual transcripts. The minimum number of clusters was used at each locus such that at least 90% of the within-locus transcript variation was retained. The processed Stringtie transcriptomes were converted to fasta format using GFFread v0.9.9 for in silico translation, for use in ortholog identification and to provide transcriptome databases for the proteomics pipeline. The Trinity-assembled transcriptomes also required processing to remove poorly supported contigs. The quality of the Trinity-assembled transcriptomes and individual transcripts within these was assessed using Transrate v1.0.3 ^73^ and BUSCOv3 ^74^. Transrate produces an overall assembly score based on the proportion of reads that provide support for the assembly and the individual contig scores. Contig scores depend on the level of read support for individual contigs. Two Transrate score thresholds were used to remove low-quality transcripts from the assemblies. A variable threshold was used to produce transcriptomes with optimal Transrate scores, referred to as strongly filtered transcriptomes. A lower threshold of 0.01 was also used to produce the weakly filtered transcriptomes. BUSCO v3 was used before and after filtering by Transrate score to assess the completeness of transcriptomes. This was done by testing for the presence of single copy orthologs that are universal within the metazoa.

### Ortholog Identification

Pairs of orthologous transcripts between the two species for each method and tissue type were identified using the reciprocal best hits (RBH) method for use in PCA and differential expression analysis. The ON transcript sequences for each tissue type and method were mapped to their respective PN transcriptomes and vice versa using blastn v2.7.1+ ^54^. Transcript pairs that were each other’s’ highest scoring match were identified as orthologs.

### Species-Specific Transcript Identification

To identify transcripts that were only expressed in one species or the other, assembled transcripts from ON were compared to those from PN and vice versa using blastn v2.7.1+ Transcripts were classed as species-specific if they did not have a match in the opposing species with at least 80% identity.

### Identification of Novel Species Specific Translation Products

Species-specific transcripts were compared to the reference annotations for their respective species using GFFcompare v0.10.1. to identify species-specific intergenic and intronic transcripts. If these transcripts had evidence of translation then the resulting translation products were classed as species-specific novel translation products.

### Principal Component Analysis

Principal component analysis (PCA) was performed in R to separate samples based on the expression of orthologous transcripts. For Stringtie WR and NR expression values were in fragments per kilobase of transcript per million mapped reads (FPKM) and for Trinity expression values were count data for equivalent orthologous transcript sections (explained in more detail below). PCA was also used to separate samples based on the expression values of orthologous proteins.

### Differential Transcript Expression Analysis

Differential expression analysis was carried out to compare the expression levels of orthologous transcripts between species and to ascertain whether expression levels had diverged more in the liver or in the testes.

Differential expression analysis for Stringtie-assembled transcriptomes was performed using a custom R script based on the Ballgown Bioconductor package. Sample FPKM values were log_2_ transformed and normalised for library size using a 75th percentile normalisation. Linear models were constructed for each pair of orthologous transcripts to predict expression levels either including or excluding species as a predictor variable. The abilities of the two models to explain the normalised expression values were compared using F-tests, with Benjamini-Hochberg multiple testing correction. Expression levels were compared between species for both liver and testes.

For Trinity differential expression analysis, orthologous pairs of transcripts were truncated to remove non-corresponding transcript sections based on the blastn mapping of orthologous transcripts to each other. This was done to account for the large differences in length found between some orthologous transcript pairs. Counts for the truncated transcripts were estimated using RSEM v1.2.31 ^75^. Differential expression analysis was carried out on the truncated-transcript count data using generalised linear model quasi-likelihood F tests in EdgeR with Benjamini-Hochberg multiple testing correction.

### Gene Ontology Annotation

GO annotation was used to assign putative biological functions to the DE transcripts and species-specific transcripts. Amino acid sequences for these proteins were predicted from the longest open reading frames of their transcripts using Virtual Ribosome v2.0 ^76^. The amino acid sequences were analysed using InterProScan 67.0 to identify families, domains and important sites and assign GO annotations ^77^. GO annotations were visualised with Blast2GO v5.0 ^78^.

### Comparison of Transcriptome Assembly Methods

For each stage of the data analysis the results found using each of Stringtie WR, Stringtie NR and Trinity were compared to find the overlap in the transcripts identified as differentially expressed or species specific. The Trinity assembled transcriptomes were mapped to their respective genomes using GMAP version 2017-11-15 to provide the genomic coordinates of the transcripts. The matching transcripts present in the transcriptomes assembled using each of the three methods were identified using GFFcompare v0.10.1.

### Substitution rate calculations using phyloP

phyloP from Phylogenetic Analysis with Space/Time Models (PHAST) v1.5 package, was used to identify the genomic sequences that evolve with a rate different than that expected at neutral drift ^79 34^. First, a neutral substitution model was constructed using phyloFit in PHAST by fitting a time reversible substitution ‘REV’ model on the phylogeny obtained from four-fold degenerate (4D) sites **(Supplementary figure 4)**. This phylogeny has topology and branch lengths similar to the subtree similarly constructed by Brawand et al using 4D sites from alignment of 9 teleost genomes (which includes the 5 cichlids genomes that we have used) ^33^ The 4D sites were extracted using msa_view from PHAST based on ON’s protein coding sequences. The five-way whole genome alignment of *O. niloticus*, *N. brichardi*, *A. burtoni*, *M. zebra* and *P. nyererei* genomes and ON’s annotation file provided by Brawand et al ^33^ was used in this analysis.

phyloP was then applied using the likelihood ratio test (LRT) method and an ‘all branches’ test to predict conservation-acceleration (CONACC) score for every site in the whole genome alignment. The output of phylop was stored in fixed-step wig format. The wig files were then converted into bed format for further analysis using wig2bed function in BEDOPS v2.4.35 ^80^. The calculated score was then mapped on the ON’s different annotation features like CDS, exons, introns, 5’UTR, 3’UTR, intergenes, ancestral repeats (AR) and novel regions using bedmap and bedops functions from BEDOPS. We compared the distributions of CONACC scores for different features and compared them using Welch t-test in R v3.6.0. As the number of novel regions were very few compared to AR; sampling from AR regions was done 10000 times, to pick per novel region; an AR which was equi-sized to the novel transcript.

Before predicting the scores, the five-way whole genome multiple alignments (mafs) were first filtered using mafFilter ^81^ to discard blocks which have sequences less than five and to remove gap only columns from the blocks. The filtered mafs were then sorted using ‘maf-sort.sh’ script from LAST (https://github.com/UCSantaCruzComputationalGenomicsLab/last.git) ^82^

Broad annotations for CDS, exons, introns and UTRs of ON were downloaded in BED format from Cambridge cichlid browser (http://em-x1.gurdon.cam.ac.uk/cgi-bin/hgTables?hgsid=21982&clade=vertebrate&org=O.+niloticus&db=on11&hgta_group=genes&hgta_track=rmsk&hgta_table=0&hgta_regionType=genome&position=LG2%3A1959784-2269783&hgta_outputType=bed&hgta_outFileName=).

Intergenic regions were assumed to be the regions that are not annotated in the whole genome and were identified by using bedtools complement (-i WholeGene.bed -g file_having_chromosome_sizes) ^83^. Ancestral repeats (ARs) were defined to be repeat masked sequences from ON that are also conserved in teleosts. The AR regions were downloaded from cichlid genome browser by taking an intersection (having at least 80% overlap) between repeat masked regions from ON and 8-way cichlids multiple alignments (On_Mz_Pn_Ab_Nb_oryLat2_gasAcu1_danRer7_maf). The annotation for all these features were downloaded for the O. niloticus assembly: Broad oreNil1.1/oreNil2.

### Divergence time calculation

Divergence time between *O. niloticus* and *P. nyererei* based on the nORF regions was carried out by using the Bayesian Evolutionary Analysis Sampling Trees model (BEAST) ^35^, which was run on BEAST v1.10.4 ^55^. The settings used in the programme were based on those used by Meyer et al. ^84^ and are as follows. Sequence evolution was taken to follow the HKY model ^85^ and the species-tree prior was set to the Yule speciation process ^86^ (Yule 1925). Empirical base frequencies were used and no site heterogeneity was assumed. A strict molecular clock was set, to allow for the most reliable comparison between trees based of nORFs sequences. The sequences were extracted from the Cambridge Cichlid Genome browser by specifying the nORF and AR sequence coordinates in *O. niloticus* (Broad OreNil1.1/OreNil2 assembly), and extracting the orthologous sequence of the other Cichlid species using the 8-way comparative genomic track option. The DNMT1 gene sequence was extracted manually from NCBI for all the species. The molecular clock was time calibrated with a fossil time constraint. The constraint was set as a lognormal prior distribution with a mean in real space of 45.5 million years ago (MYA) and a standard deviation of 0.5 MYA. This time calibration was based on cichlid fossils estimated to be 45 million years old ^56^. The substitution rate was fixed to allow better comparison between trees. The neutral model was set at 1 and any deviation from this was taken into account while building the trees. A chain length of 10 million Markov Chain Monte Carlo (MCMC) was used to construct each tree.

